# Mechanoepigenetic Targeting of Histone Methyltransferase EZH2 Increases Potential of Scalable Manufacturing in Human Derived Primary Mesenchymal Stromal Cells

**DOI:** 10.64898/2025.12.10.693526

**Authors:** Lauren Monroe, Samantha Kaonis, Soham Ghosh

## Abstract

Manufacturing clinical grade mesenchymal stromal cells (MSCs) remains a major bottleneck for cell-based therapies, as extensive in-vitro expansion on standard tissue-culture plastic (TCP) drives loss of stemness, reduced immunomodulatory activity, and diminished therapeutic efficacy. Although substrate stiffness is known to influence MSC fate through mechanotransduction, the epigenetic mechanisms linking mechanical stress to progressive phenotypic drift remain poorly defined. A mechano-epigenetic pathway is identified in this work, centered on the histone methyltransferase EZH2 - that governs chromatin remodeling and loss of stemness during serial passaging. Multi-omics, high-resolution imaging, and functional assays show that MSCs expanded on mechanically stiff TCP accumulate H3K27me3 repressive chromatin mark, lose SWI/SNF -ARID1A chromatin-remodeling foci, and exhibit an altered chromatin accessibility profile.

Pharmacological inhibition of EZH2 with GSK343 selectively reduced H3K27me3, restored ARID1A-containing SWI/SNF organization, and preserved MSC morphology and expression of canonical stemness markers (CD73, CD90, CD105) even at later passage. ATAC-seq analysis revealed that GSK343 rebalanced chromatin accessibility, reopening TEAD/YAP-responsive regulatory regions while repressing accessibility at lineage-priming and senescence-associated sites. RNA-seq demonstrated that GSK343 maintained transcriptional programs associated with immunomodulation, migration, and trophic signaling, while suppressing hyperproliferative and senescence-associated genes that are characteristic of late-passage MSCs. Proteomic profiling of MSC secretomes further showed that GSK343 attenuated pro-fibrotic ECM factors and senescence-linked proteins while enhancing angiogenic and reparative mediators. Functionally, conditioned media from GSK343-treated MSCs significantly increased primary chondrocyte proliferation, demonstrating preserved therapeutic potency.

Together, these findings establish EZH2 as a central mediator of stiffness-induced epigenetic drift in human MSCs and demonstrate that EZH2 inhibition can maintain the stemness without compromising their expansion ability. This work provides a foundational strategy for mechano-epigenetic engineering of MSCs and highlights EZH2 inhibition as a scalable, manufacturing-compatible approach to preserve potency for regenerative medicine applications.

**SIGNIFIANCE STATEMENT:** Primary mesenchymal stem/ stromal cells (MSC) derived from bone marrow are currently under numerous clinical trials for regenerative medicine and cancer treatment. Despite showing tremendous potential for their differentiation, trophic and immunomodulatory properties, their efficacy in clinical endpoint is limited. The traditional cell culture condition in cell culture flask limits their clinical translation by changing the MSC phenotype with serial passaging, a critical step required for harvesting a clinically appropriate number of cells. This work discovered the precise epigenetic mechanism involving histone methylation responsible for such changes in MSC phenotype. By inhibiting the histone methylation by GSK343, this study showed that MSC phenotype can be maintained over serial passaging as demonstrated by microscopy, multi-omics, and in vitro functional assays.

## INTRODUCTION

Mesenchymal stem/stromal cells (MSCs) are widely used in a myriad of regenerative medicine and immunoengineering therapy applications due to their multipotency, immunomodulatory abilities, ant- inflammatory and trophic properties [1,2]. Hundreds of clinical trials are currently using MSCs for various applications to treat musculoskeletal. autoimmune, cardiovascular, neurological and cardiovascular diseases [3]. However, despite their broad use for decades, their clinical efficacy remains inconsistent.

For MSCs to be used in cell based or acellular therapies, they must be expanded in vitro over multiple passages to achieve a large number on the order of hundreds of millions of cells. During this expansion process, MSCs lose their innate properties including their multipotency and self-renewal capabilities and an altered secretome due to their phenotypical changes. This shift severely limits their clinical outcomes. However, there is a lack of a clear mechanistic understanding of why MSCs lose their original early passage specific phenotype during expansion on conventional tissue culture plastic (TCP). MSC fate is highly subjected to the mechanical properties of their surrounding environment, such as their substrate or scaffold stiffness. Cells can sense the mechanical property by mechanotransduction and can transmit information to the nucleus to regulate a wide range of cellular processes including epigenetic regulations that can change the gene expression profile. MSCs display a spindle-like morphology on a softer substrate that mimics the bone marrow stiffness and on a stiffer substrate they show a flat, stress fiber reach morphology [4]. Standard TCP is orders of magnitude stiffer (∼GPa range) than native MSC niches (∼kPa range). A mechanically stiffer environment has shown to affect cellular architecture such as increasing cytoskeleton tension, higher nuclear deformation, upregulating lamin A/C production, and inducing global chromatin remodeling [5, 6, 7, 8, 9, 10, 11, 12, 13, 14].

In MSCs, matrix stiffening has been shown to cause an increase in histone acetylation by decreased HDAC activity [13]. Additionally, soft substrates promote chromatin enrichment at the nucleus periphery where stiffer substrates have a more even distribution throughout the cell’s nucleus [15]. This indicates that chromatin undergoes spatial reorganization based on substrate stiffness [15]. H3K4me3, an active euchromatin mark, was found to be decreased in response to softer substrates and conversely, H3K27me3 was enhanced on soft substrates, particularly at the nuclear periphery, indicating increased levels of chromatin condensation on soft compared to stiff substrates [15].

Previous and emerging work has also suggested that stem cells not only respond to current mechanical cues but also retain their mechanical memory through persistent epigenetic marks. MSCs have been shown to have an increase in acetylation levels on a stiff substrate; however, if MSCs are only seeded on a stiff substrate for a short time and then the substrate is softened, rapid chromatin changes occur including the transport of YAP out of the nucleus, an increase in chromatin condensation, and ultimately a decrease in acetylation levels. On the other hand, if the MSCs are seeded for a longer time on a stiff substrate, the epigenetic profile is irreversible [12, 16, 17]. Overall, the substrate stiffness changes induce long-lasting chromatin modification that can influence future differentiation potential and transcriptional activity even after the mechanical environment is altered. Overall, studies indicate that soft substrates help preserve the native MSC phenotype; unfortunately, they do not fully resolve the problem, nor do they scale well for manufacturing purposes.

Since studies suggest that chromatin level changes likely drive this loss of stemness, this makes epigenetic regulation an appealing target. Many epigenetic drugs have been validated to block or modulate histone modifications such as methylation and acetylation. However, it remains unclear which specific epigenetic features change during MSC expansion on stiff TCP and if pharmacological targeting of MSC epigenetic landscape can be a solution to maintaining the MSC phenotype during serial passaging.

The goal of this study was to identify key epigenetic alterations that occur during serial passaging on stiff (TCP) and soft (bone marrow-like) substrates and to determine whether targeted epigenetic modulation can preserve MSC stemness and its functional potency. To achieve this, immunofluorescence staining of key epigenetic marks was performed to map the overall chromatin architecture during serial passaging. Drugs to target epigenetic modifiers such as EZH2 and KDM4 were evaluated alongside cells passaged on stiff and soft substrates to determine whether they helped restore or preserve a higher stemness phenotype. ATAC seq, RNA seq, and immunofluorescence were used to investigate the mechanisms of chromatin level changes, and its restoration by epigenetic modifications. The resulting MSC populations were evaluated using RNA-seq, to analyze the transcriptional stemness programs, proteomics to investigate their secretome, and a functional assay of chondrocyte proliferation to inspect their therapeutic potency. By investigation of the epigenetic alterations during serial passaging, as well as demonstrating that targeted epigenetic modulation can reverse this, this study provides a mechanistically grounded strategy to possibly unlock clinically scalable MSC manufacturing.

## MATERIALS AND METHODS

### MSC culture

Passage 2 (P2) primary human bone marrow derived MSC (hBM-MSC) was purchased from Lonza for all experiments. These cells are derived from a single donor, a healthy adult individual. Cells were maintained in bulletkit (Lonza) growth medium kit consisting of MSC Basal medium (PT-3238) and the supplemental kit (PT- 4105) as stated in the manufacturer’s protocol. The culture condition was 5% CO2, 37°C temperature and 90% relative humidity. P2 cells were thawed and immediately seeded in a 6-well plate, 3 wells per group, and passaged at 90% confluency with a starting density 5000 cells/ cm^2^ to P5. Trypsin-EDTA (0.05%) was used for subculturing. Media was changed 24 hrs after seeding and every 2-3 days.

### MSC culture on substrate with different mechanical stiffnesses

Two different substrate stiffness groups were prepared for cell culture. Soft: Sylgard 527 PDMS (*E*∼5 kPa) and Stiff/Control: tissue culture plastic (*E*∼1 GPa). These mechanical stiffness conditions represent the physiological bone marrow stiffness and the standard T flask cell culture conditions respectively. Untreated polymer bottom slides or plastics were used for the stiff group. For the soft substrate group, the 527 PDMS was cast directly in the wells of the 6-well plate as described in section “Preparation of PDMS substrate”. P3 and P5 cells were also seeded in either soft of stiff 8-well slides (ibidi, 80821) used for immunostaining and imaging. For the soft group 8-well slide, the bottom polymer coverslip was removed, and custom-made chamber slides were refabricated with a thin PDMS substrate as described in section “Preparation of PDMS substrate”. Treated 8-well plates (ibidi, 80821) were used for the stiff group. Some cells were harvested by trypsinization from each group at P3 and P5 for RNA and ATAC sequencing.

### Cell treatments with epigenetically modifying drugs

For all drug treatment groups, the drugs were applied to their respective groups 24 hrs post seeding after each new passage from passage 3 (P3) to passage 5 (P5). GSK126 (5 μM) was used to block epigenetic modifier EZH2 which is the catalytic enzyme within the Polycomb Repressive Complex 2 (PRC2) responsible for tri- methylating H3K27me3. GSK343 (2 μM and 1 μM) was also used to block epigenetic modifier EZH2; however, it has been shown to be less selective for EZH2 over EZH1 compared to GS126. ML324 (1 μM) was used to block epigenetic modifier KDM4 of H3K9me3. All non-treated groups also received a media change at 24 hrs post seeding.

### Preparation of PDMS substrate

Polydimethylsiloxane (PDMS) substrate was prepared using Sylgard 527 (Dow Corning). To fabricate the thin soft PDMS substrate (∼ 5 kPa), Sylgard 527 was mixed at a 1:1 ratio of each solution provided by the manufacturer and vigorously mixing. The mixture was then placed in a water bath at 37°C for 20-30 minutes to begin the curing process. For coating the 6-well plates, the mixture was poured in the center of each well (∼2 mL) and the plate was rotated around to spread the mixture across the whole well surface and placed in an oven at 75°C to cure overnight. For the 8-well thin gel preparation, a small amount of the mixture (∼70 μl) was distributed across the top of a glass coverslip (24x60 mm). The glass coverslips were then placed in a vacuum chamber for 20 minutes to remove any bubbles. For attachment of the coverslip to the bottomless chamber slide (ibidi, 80821), the coverslip was partially cured for 2 hours at 75°C. Sylgard 184 (Dow Corning) PDMS sample was prepared by mixing ten parts of base with one part of curing agent and vigorous mixing. This mixture (Sylgard 184) was then used to mount the coverslip to the chamber slide. To completely cure, it was placed in a 75°C oven overnight. Cured PDMS were sterilized in the cell culture cabinet with a 70% ethanol soak under UV light for 20 minutes. The plates were then plasma treated (BD-20AC, Electro-Technic) and incubated for an hour at room temperature in a type I collagen coating solution (50 μg/ml, A10644-01, Gibco) to improve cell attachment and proliferation. Cells were seeded on these prepared substrates and cultured with the previously stated culture conditions.

### Immunofluorescence staining and imaging

For all imaging tasks, cells were fixed with 4% PFA (in 1X PBS: Phosphate Buffer Solution) for 7 minutes at room temperature. Cells were permeabilized with 0.1% Triton X-100 in 1X PBS, washed with 1X PBS, blocked for the non-specific binding sites with bovine serum albumin and normal goat serum. Primary antibodies were diluted in a buffer and incubated overnight at 4°C. Next, secondary antibodies were diluted and added at room temperature for 2 hours. DNA was stained using DAPI (ThermoFisher Scientific). Cells were imaged across their midsection using a confocal microscope (Zeiss LSM 980 with Airyscan 2). Cells were imaged either at low resolution with 10X air objective (confocal microscopy) or at high resolution with 63X oil objective (confocal or super resolution) for all subsequent analysis. For all imaging, the imaging setting (laser power, gain etc.) was consistent between the groups. List of dilution of primary antibodies (all from Invitrogen): CD73 – 1:100, CD105 – 1:100, VEGF-1:100, CD90 – 1:100, H3K9me3 – 1:500, H3K9ac – 1:400, H3K27me3 – 1:500, ARID1A – 1:400, Lamin – 1:400. All secondary antibodies – either anti-rabbit or anti-goat (Invitrogen) were used at 1:200 dilution.

### Quantification of immunofluorescence images

For quantification of functional marker intensities (CD73, CD90, CD105, and VEGF), cell profiler was used to measure the average intensity of pixels per cell and the average intensity of pixels in the nucleus from the DAPI stain. The intensity of the functional marker per cell was then normalized to the DAPI intensity of that cell to get a normalized intensity value of the functional stain per cell. For quantification of epigenetic marker intensities (H3K9me3, H3K27me3, and H3K9ac), cell profiler was used to measure the average intensity of pixels in the nucleus by using the DAPI channel as a mask. The DAPI intensity per nuclei was also measured and used to get the normalized values of the average intensity per cell as previously stated. To quantify the distribution of a stain within the nucleus from the super resolution images (such as for the epigenetic marks, ARID1A, and lamin stains), ImageJ was used to identify the nuclear boundary and measure the skewness and kurtosis values of the desired stain per nuclei.

### RNA sequencing and analysis

Designated cells from all treatment groups were collected by trypsinization and then spun down into a pellet. Total RNA was extracted from cells using the RNeasy MiniKit (Qiagen) according to the manufacturer’s protocol. RNA quantity and quality were measured on the NanoDrop One Spectrophotometer (ThermoFisher). RNA samples were then sent to Novogene for sequencing. Analysis of RNA sequencing was done in the Partek software (Partek Inc., St. Louis, MO). The raw sequence reads were first subjected to quality control checks and then aligned to the human reference genome (hg38) using STAR alignment. Quality checks were performed on aligned data and HTseq was used to count aligned reads. Gene counts were filtered to filter out gene counts of 0 in at least 80% of the groups and median ratio was used to normalize the count data. Lastly, DESeq2 was used for differential expression analysis to identify any differentially expressed genes between specific sample groups.

### ATAC seq and analysis

Cells were cultured and treated according to the same protocol found in the “Cell culture and pharmacological treatment” section. Approximately half million cells per sample were harvested, washed in PBS, and snap-frozen in MSC freeze media (Lonza) before shipping to a commercial vendor (Novogene) for ATAC-seq library preparation (using the Omni-ATAC protocol with Tn5 transposase) and paired-end sequencing (2×150bp, ∼50M reads per sample). Quality control parameter for sequencing was set at the total cell amount >50000 and cell viability of >60%, the standard practice by Novogene. Raw FASTQ files were assessed using FASTQC (v0.12.1) and adapters/low-quality bases were trimmed with Cutadapt (v2.6). Trimmed reads were aligned to the GRCh38 human genome using Bowtie2 (v2.3.5.1) and duplicated. Mitochondrial reads were excluded to reduce noise. Open chromatin regions were identified using MACS2 (v2.2.6; parameters: default settings and --nomodel) in paired-end mode. Bedtools (v2.30) was used to quantify the number of strictly shared peaks and unique peaks between treatment groups to generate the Venn diagram. Downstream analysis and normalization were performed using counts per million (CPM). Differential accessibility was assessed by comparing |FC| > 2 and an average normalized count >10 due to the lack of biological replicates. To examine transcription factor (TF) binding motifs associated with significant changes in chromatin accessibility, motif enrichment analysis was performed using HOMER. v4.11 suite (Hypergeometric Optimization of Motif EnRichment) (Heinz et al., 2010). Peak sets, one for significantly upregulated peaks and one for significantly downregulated peaks, were converted to BED format and supplied to HOMER as input regions for motif discovery. Only motifs meeting the significance cutoffs (p<0.05) and exhibiting a match score of at least 70% to a known TF family were considered for interpretation.

### Mass Spectrometry and analysis

Media from cultured MSCs at passage 5 from Control, Soft, GSK126, and GSK343 2 µM groups was collected, lyophilized, and sent to the CSU ARC-BIO facility for mass spectrometry analysis. Lyophilized samples were reconstituted, sonicated, and subjected to overnight acetone precipitation, followed by centrifugation, acetone washing, air-drying, and resuspension in lysis buffer. Protein concentrations were determined using the Pierce BCA assay. For digestion, 100 µg of protein per sample was reduced, alkylated, and digested with Trypsin/Lys-C using the EasyPep Mini MS Sample Prep Kit; peptides were cleaned, dried, and resuspended in 3% acetonitrile/0.1% formic acid, and concentrations were estimated by A205. Approximately 1 µg of peptides was separated by reverse-phase nanoLC (Vanquish Neo) using a 90-min gradient and analyzed on an Orbitrap Eclipse mass spectrometer with MS1 acquisition at 240,000 resolution, HCD MS/MS (NCE 30%), and dynamic exclusion. Data were processed in Proteome Discoverer 3.0 using Sequest HT against the human reference proteome with cRAP contaminants, with carbamidomethylation (static) and methionine oxidation (dynamic) specified, and precursor/fragment tolerances of 10 ppm and 0.6 Da. INFERYS rescoring and Percolator were applied to achieve an FDR ≤1%, and peptide abundances were normalized across samples by total ion intensity (<u>cite</u>). Fold change analysis was conducted among the ratios of samples GSK126, GSK343, and Soft over the Control sample. No statistical tests were performed due to only one replicate per condition being tested. Higher abundance proteins or significant levels of proteins were defined if |FC| was greater than 2.

### Chondrocyte proliferation assay

MSC conditioned media (CM) from treatment groups Control P5 and GSK343 2 µM was collected and frozen at 20C in aliquots. Primary rabbit chondrocytes were cultured in MSC CM in a 1:1 and 3:1 ratio of basal media to MSC CM. Control wells consisted of the same ratios but with MSC media instead of MSC CM.

### Statistical Analysis

Statistical analysis was performed using a post-hoc Tukey’s test in RStudio. Results are expressed as violin plots or bar graphs with mean and standard deviation as error bars, with p values specified in the figure or figure legends. Sample and replicate numbers are provided in the figure legends.

## RESULTS AND DISCUSSIONS

### EZH2 inhibition by GSK343 maintains the MSC phenotype in later passage emulating the culture on a soft substrate

A softer substrate resembling bone marrow stiffness is known to better maintain the MSC phenotype. Previous studies have shown that MSCs serially passaged on softer substrates maintain their morphology and proliferative ability [18]. Additionally, MSC specific markers show a higher expression after culture on soft substrates compared to stiffer substrates [19]. Epigenetic targeting was performed to investigate if those treatments can emulate the MSC specific phenotype at a later passage. GSK126 and GSK343 were used to inhibit the histone methyltransferase EZH2, that catalyzes H3K27 tri-methylation, and ML324 was used to inhibit JMJD2, a histone demethylase known to demethylate H3K9me3 [20, 21, 22].

Using immunofluorescence (IF), the relative expression of key MSC surface markers (CD73, CD90, CD105) and the functional marker VEGF was quantified across different substrate stiffnesses (soft substrate, E ∼ 5 kPa and cell culture plastic, E ∼ 1GPa) and epigenetic treatments over multiple passages (passage 3 to 5) (Fig. 1). The passages were chosen from the well-established clinical bottleneck where MSC lost their phenotype from passage 3 to 5. MSC on a soft substrate showed a significant increase in CD73 expression levels compared to both early passage (P3) groups and the Control (plastic) P5 group. All the epigenetic interventions also increased the CD73 expression levels as shown by an increase in mean intensity compared to control and soft groups. Similarly, CD105 expression indicated the same general trend. GSK343 treatments at different concentrations and the Soft P5 group were shown to have significantly increased levels of CD105 expression compared to all other groups. However, ML324 and GSK126 drug groups showed significantly less expression than both GSK343 groups and were more on par with the Control P5 and P3 groups.

**Figure 1.**
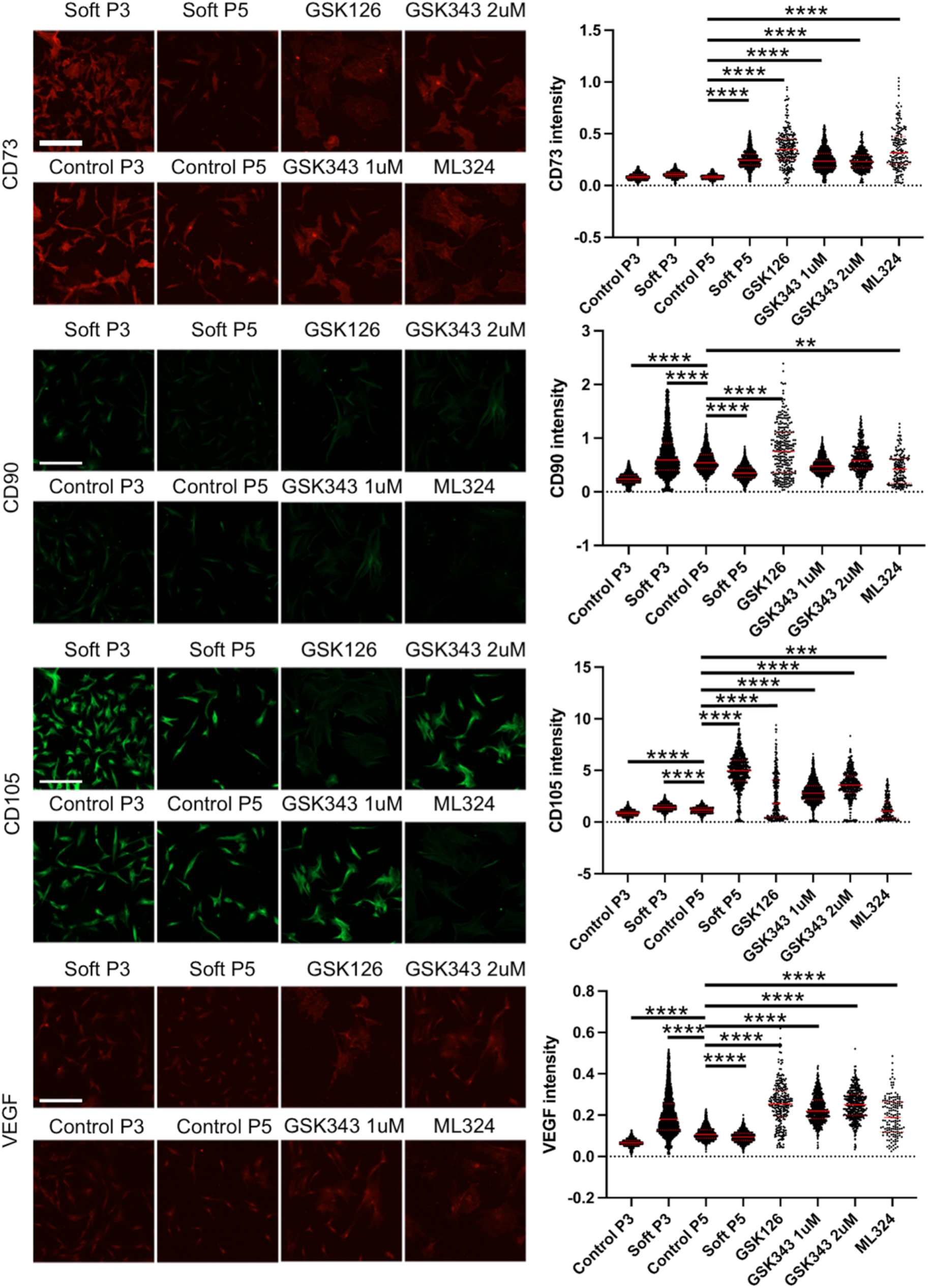
Effect of substrate stiffness and epigenetic priming on primary hBM-MSC specific markers. Representative images and IF intensity analysis of MSC specific surface markers including CD73, CD90, and CD105 and MSC functional marker VEGF. (n>200, ****p<0.0001, ***p<0.001). Scale bar: 300μm. GSK343 at 2µM concentration exhibits an overall better maintenance of MSC phenotype, similar to a soft substrate by increasing key MSC surface and functional marks.

CD90 intensity appeared more variable. GSK126 and Soft P3 treatments show the highest intensity levels with GSK343 2 μM and Control P5 groups trailing behind them. Additionally, Control P3 continues to have significantly lower expression levels than the others. Compared to Control P5 groups, none of the drug treatments resulted in a higher level of CD90, but GSK343 at 2 μM maintained the same level of CD90 compared to Control P5 and a higher level compared to Soft P5.

The functional marker VEGF expression levels also generally followed the same trend as was in the CD73 and CD105 MSC surface markers. Both GSK343 concentrated groups and GSK126 treatment had significantly higher mean intensities than all other groups. ML324 and Soft P3 groups were also significantly higher than the untreated P5 and Control P3 groups.

This intensity analysis suggests that GSK343, the EZH2 inhibitor emulates the culture on a soft substrate and better maintains the MSC stemness based on the most critical CD markers, compared to the late passage cells cultured on plastic.

### The MSC epigenetic signature and chromatin architecture is altered by serial passaging and epigenetic modifications

Histone modifications such as methylation and acetylation are recognized to play an important role in MSC fate. Specifically, dysregulation of histone modifications can lead to changes in their transcriptome profile, lineage commitment, and overall phenotype [23, 24, 25]. For this study, heterochromatin marks H3K37me3 and H3K9me3 indicative of condensed chromatin and gene repression along with euchromatin mark H3K9ac showing open chromatin regions and active gene transcription were investigated. Immunofluorescence intensity of these epigenetic modifiers was analyzed in MSC nuclei at early and late passages from the same treatment groups as above to study the global level chromatin status in MSC (Fig. 2).

**Figure 2.**
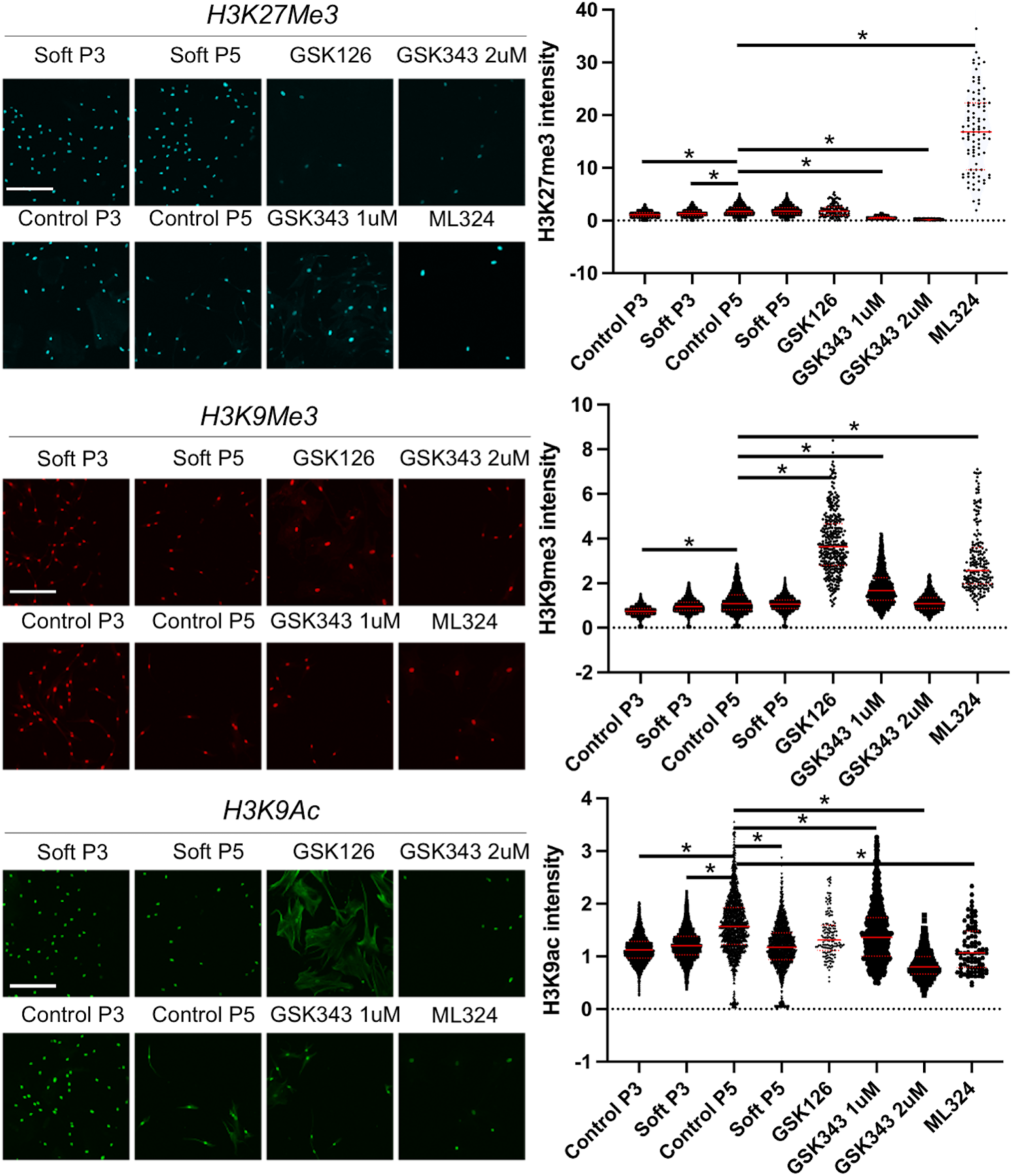
Effect of substrate stiffness and epigenetic priming on key histone modification marks. Representative images and IF intensity analysis of epigenetic markers including H3K9me3, H3K27me3, and H3K9ac. (n>200, *p<0.0001). Scale bar: 300μm. Epigenetic modifiers significantly change the epigenetic landscape by changing the degree of chromatin condensation inside the MSC nucleus.

H3K27me3 is a hallmark of facultative heterochromatin meaning that it can switch between condensed and open chromatin states depending on its environment and transcriptional need. GSK343 inhibits EZH2 therefore lowering H3K27 methylation. H3K27me3 intensity levels were significantly lowered by GSK343 treatment indicating that the drug successfully blocked epigenetic modifier EZH2 causing a decrease in H3K27me3 expression. Additionally, a general trend showing higher H3K27me3 expression levels in P5 than P3 cells suggests an overall chromatin compaction with passaging irrespective of substrate stiffness. ML324 treated group displayed a significantly higher mean intensity across all treatments, suggesting significant gene repression where GSK126 treatment showed a mean value similar to the P5 untreated groups.

The intensity analysis of H3K9me3, a constitutive heterochromatin marker denoting permanently packed or condensed chromatin, showed the largest increase in ML324 and GSK126 mean intensity suggesting a global conversion of euchromatin to heterochromatin. GSK343 1 µM treatment also showed a significantly higher H3K9me3 intensity; however, GSK343 2 µM treatment showed no significant difference in mean intensity compared to the Control P5 group lending to the conclusion that H3K9me3 isn’t largely affected by 2 µM GSK343 intervention. Additionally, the Control P3 group showed an overall lower level of constitutive heterochromatin indicating chromatin condensation has a positive correlation with passaging, and GSK343 at 2 µM maintains the status quo of H3K9Me3 compared to the early passage level.

Euchromatin mark H3K9ac expression level was highly variable across treatments. Control P5 showed higher expression levels than both P3 groups. However, the Soft P5 group had a lower intensity compared to the Control P5 groups that had an intensity level closer to that of the P3 groups indicating that passing on a soft substrate may maintain but not increase additional open chromatin regions. All epigenetically modified groups showed lower H3K9ac expression compared to Control P5 with the GSK343 2 µM group showing the lowest H3K9ac expression. These results further demonstrate that epigenetic modifiers can change the epigenetic landscape to affect the MSC phenotype and expression profiles.

Previous studies indicated that distribution of chromatin and epigenetic modifiers inside the nucleus can be related to cellular function. For example, H3K9Me3 is homogeneously distributed in clusters in cardiac stem cells. As they differentiate into mature cardiomyocytes, H3K9me3 is localized at the periphery whereas the cardiac fibroblasts show larger H3K9Me3 foci [26]. Epigenetic modifiers can also create localized nanodomains in MSC as another study showed [15]. Therefore, super resolution imaging was performed to visualize the distribution of key epigenetic marks inside the nucleus over serial passaging (Fig. 3). To measure the chromatin distribution for the various epigenetic marks, skewness and kurtosis were calculated based on the distribution of immunofluorescent signals. Skewness measures the asymmetry in the distribution of pixel intensities where a lower positive value indicates a more even distribution, and a higher skewness would have regions of higher pixel intensity against a dimmer background. Kurtosis measures how “tailed” the intensity distribution indicating whether extreme bright/dark pixels are more frequent than in a normal distribution. A higher positive kurtosis value indicates a sharper peak suggesting a larger number of bright spots and vice versa for a lower value.

**Figure 3.**
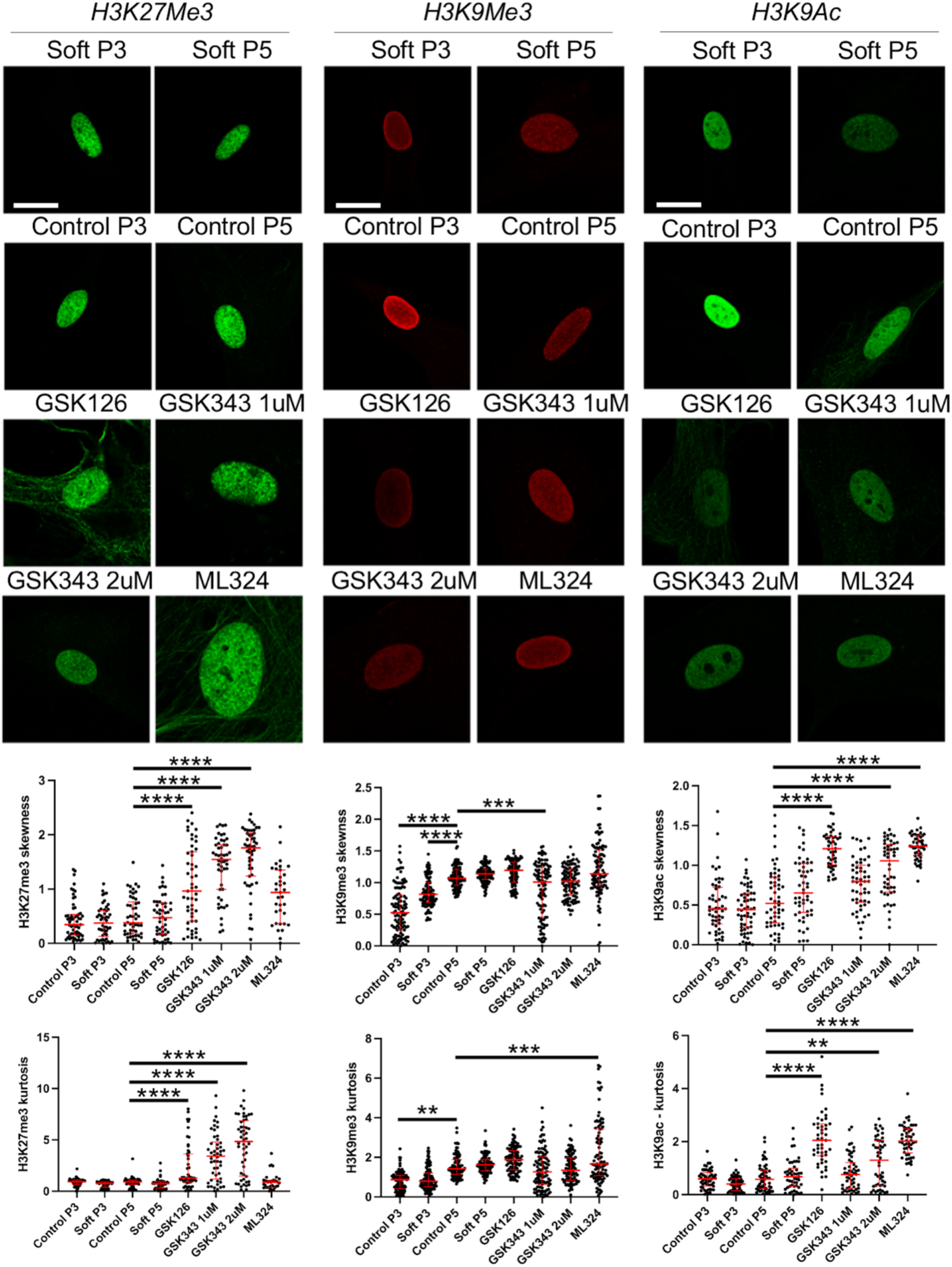
Distribution of key epigenetic marks within MSC nuclei. Representative 63X super-resolution images and quantification of intensity distribution within the nucleus through skewness and kurtosis analysis. (n=50, **p<0.01, ***p<0.001, ****p<0.0001). Scale bar: 20μm.

H3K27me3 distribution showed a significantly higher skewness and kurtosis for GSK126 and GSK343 treatments compared to the Control P5 group suggesting GSK126 and GSK343 created brighter foci and a larger number of intense foci in the nucleus where all non-treated groups showed a more homogenous distribution. H3K9me3 displayed higher skewness in Control P5 than both P3 groups. Additionally, GSK343 treatment displayed a lower skewness value than Control P5. However, kurtosis values showed that the Control P3 was significantly lower than Control P5 and GSk343 groups were not significantly different. This indicates a more heterogenous distribution of H3K9me3 in the GSK343 treated cells but not necessarily a larger number of foci. Lastly, H3K9ac distribution showed significantly higher skewness and kurtosis in all epigenetically modified groups except GSK343 1 µM compared to Control P5. This indicated that the drug treatments resulted in a greater number of localized H3K9ac foci compared to untreated groups.

### ATAC sequencing indicates EZH2 inhibition through GSK343 changes the accessibility in the MSC

MSCs seeded on a stiffer substrate have been shown to have increased acetylation levels at the later passage indicating a more open chromatin architecture [12], which matches with our data, and it is decreased by 2 µM GSK343 (Fig. 2). However, the skewness and kurtosis data (Figure 3) showed that H3K9Ac clusters in more distinct foci in 2 µM GSK343 treated group compared to Control P5. Therefore, to delve deeper into the general accessibility of MSCs seeded on stiffer substrates compared to those that received EZH2 inhibition treatment through GSK343, ATAC sequencing was performed on the Control P5 and GSK343 2 µM treatment groups.

Upon comparing strictly shared peaks and unique peaks between the GSK343 treated MSCs and control, a greater number of peaks in the Control group was evident (Fig. 4A). This indicates a more accessible chromatin architecture in the Control and that the GSK343 treatment might be causing overall chromatin compaction. Additionally, the significant differentially accessible peaks/regions (DARs) were investigated where significance was defined as average normalized counts > 10 and a |FC| > 2. This analysis also showed a similar trend with a lower number of significantly upregulated peaks (53 upregulated peaks) and a greater number of significantly downregulated peaks (78 downregulated peaks) in the GSK343 treated group versus the Control (Fig. 4B). Overall, this ATAC seq data and analysis indicates a shift in the chromatin landscape toward more chromatin compaction or gene repression due to EZH2 inhibition through GSK343 treatment but specific regions of the gene open by GSK343 and some other regions close, when compared to control.

**Figure 4.**
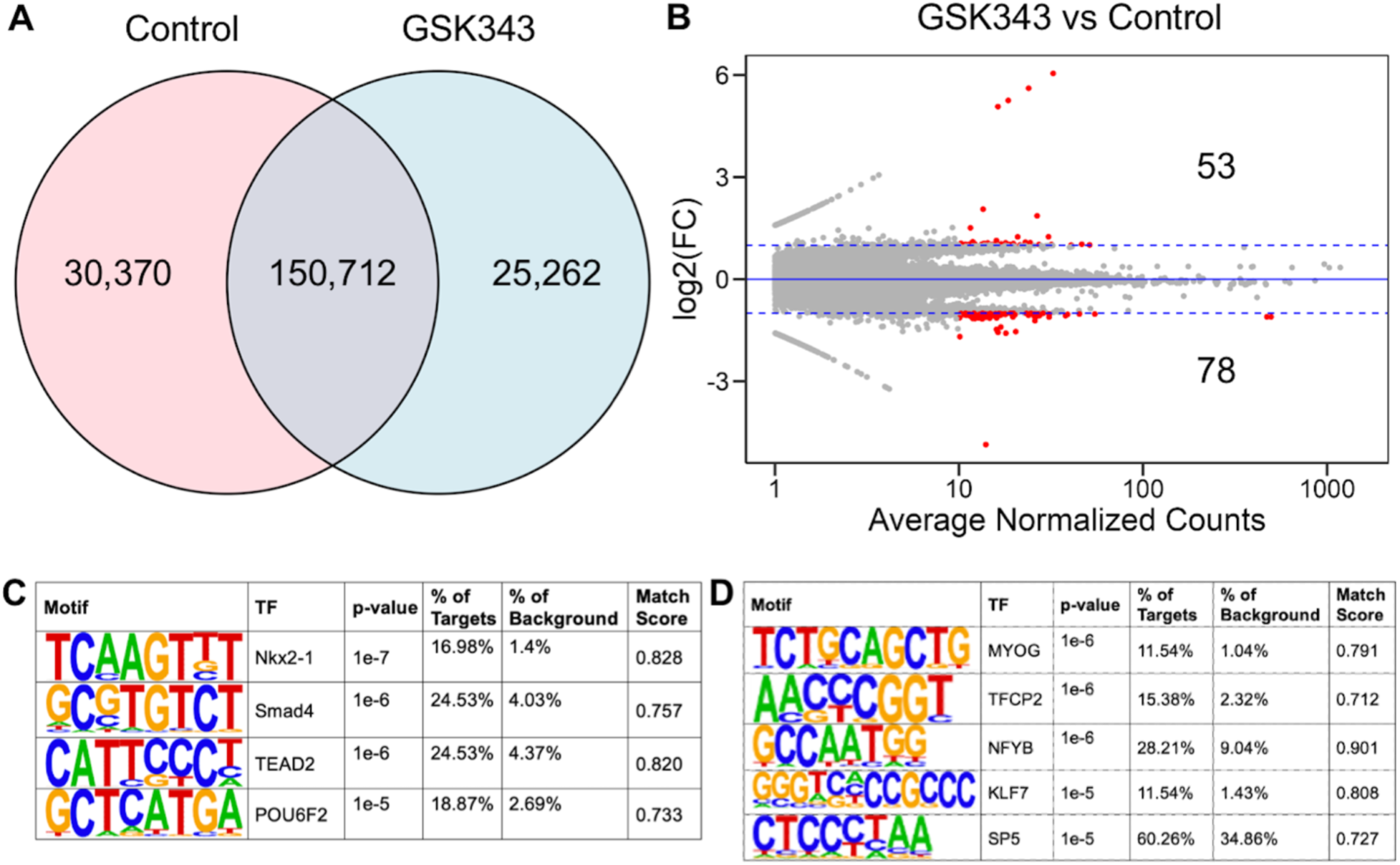
(A) Chromatin accessibility of Control and GSK343 treated MSCs from ATAC sequencing data. **(A)** Venn diagram plot showing strictly shared and unique peaks between treatment groups. **(B)** Volcano plots showing significant peaks in red. Significant peaks defined as: average normalized counts>10 and FC>abs(2). 1 replicate per treatment group was analyzed. GSK343 treatment decreases the total number of accessible peaks and shows a lower number of upregulated DARs than downregulated DARs. **(C,D) Homer *de novo* motif analysis of significant peaks with high match score. (C)** Motif results from the 53 significantly upregulated peaks in GSK343 vs Control. **(D)** Motif results from the 78 significantly downregulated peaks in the GSK343 vs Control.

Hypergeometric Optimization of Motif EnRichment (HOMER) was performed to investigate key motifs from the 53 significantly upregulated peaks and 78 significantly downregulated peaks. The Homer *de novo* motif analysis found 13 significant motifs from the upregulated peaks (Figure S1) and 12 significant motifs from the downregulated peaks (Figure S2). The most significant motifs with a match score of greater than 70% are described in Fig. 4C and Fig. S1. Notably, motifs enriched in the upregulated peak targets in GSK343 included Nkx2-1, Smad4, TEAD2, and POU6F2. Nkx2-1 and POU6F2 are markers of derepressed developmental gene programs indicating more open chromatin and more gene expression at developmental and differentiation-associated enhancers. Smad4 is a common mediator of the TGF-β and BMP pathways, both of which regulate MSC differentiation and control proliferation and senescence. An upregulation of these motifs may indicate a priming of MSCs for differentiation or alter their response to external cues. Lastly, TEAD family TFs (TEAD1/2/3/4) bind YAP/TAZ cofactors and are key regulators or mechanotransduction, proliferation, stem cell maintenance, and ECM and cytoskeletal programs. Enrichment at TEAD2 could suggest YAP/TAZ enhancers became more open, and chromatin is shifting toward a stem/progenitor-like regulatory program. Overall, enrichment at these motifs could indicate that MSCs adopted a more proliferative, remodeling, or progenitor-like state.

Motifs enriched in downregulated peaks in GSK343 group included MYOG, TFCP2, NFYB, KLF7, and SP5 (Fig. 4D and Fig. S2). This indicates that these chromatin regions are more closed, and regulatory programs associated with these TFs are reduced in the GSK343 treated group compared to the Control P5 group. MYOG is a key myogenic differentiation transcription factor indicating MSCs may be losing accessibility at muscle-lineage-associated enhancers. TFCP2 and NFYB are both involved in cell cycle regulation and proliferation, indicating a decreased accessibility at proliferation-associated promoters. KLF factors have been shown to maintain stemness programs, metabolic homeostasis, and anti-differentiation roles, among others suggesting MSCs may be losing a stem-like regulatory signature. Lastly, SP5 is a direct Wnt/β-catenin target that often acts as an early regulator of early proliferation, represses or attenuates Wnt-driven differentiation programs and is a TF maintaining undifferentiated states. Put together, these results partially oppose the motifs found from the upregulated peak targets and point to a loss of stem/progenitor state with a decrease in cell cycle and proliferative associated regulatory accessibility. Therefore, the GSK343 does not result in a linear change in stemlike character of MSC, rather depending on the specific gene the accessibility is increased in some stemness associated genes and decreased in some others.

### SWI/SNF chromatin remodeling complex is the key mediator of stiffness induced mechanotransduction in MSC that regulate the chromatin accessibility

The SWI/SNF chromatin remodeling complex (CRC) that contains ARID1A (BAF) has been shown to be functional and form distinct nuclear foci in MSCs at lower passages in a high stem-like state [27, 28, 29, 30]. Specifically, ARID1A has been shown to keep the complex poised at key stemness and lineage-control genes [31]. However, with high mechanical stress due to multiple passages at non-physiological stiffness levels these foci become more diffuse showing a more homogenous stain of ARID1A in the nuclei of late passage cells as indicated by a lower skewness and kurtosis values compared to P3 or early passage cells (Fig. 5 A, B). However, in the GSK343 treated cells, a significantly higher skewness level was found closer to the distribution of ARID1A shown in early passage cells. This points toward a dysfunctional SWI/SNF complex in late passage cells which fails to properly open the chromatin at key genomic loci. This might then create a permissive environment for the polycomb repressive complex 2 (PRC2), where EZH2 deposits the repressive H3K27me3 mark, further “locking” these regions in a closed state. Therefore, by inhibiting EZH2 through GSK343, H3K27me3 deposition is inhibited, and key genes are kept accessible to transcription factors.

**Figure 5.**
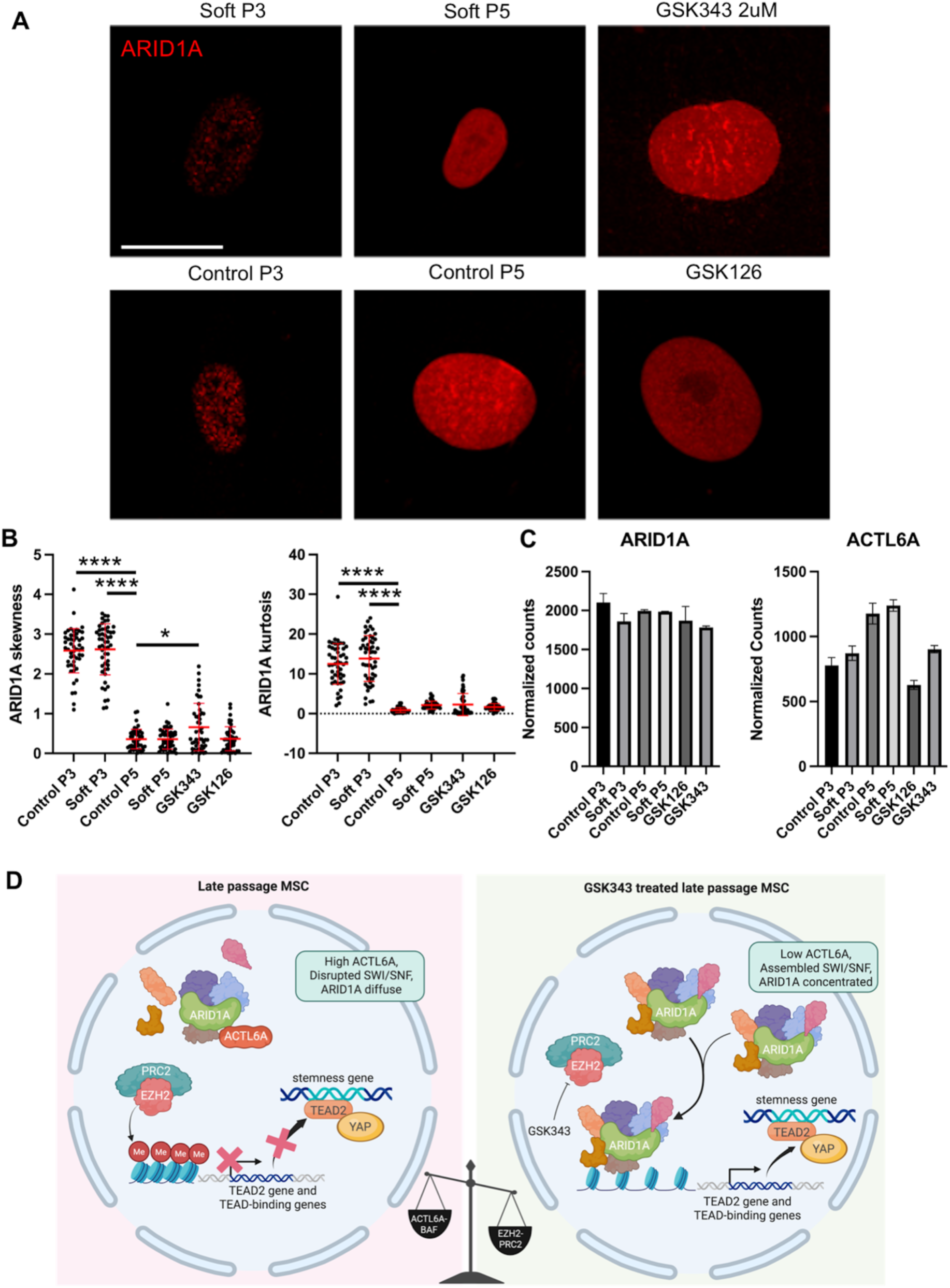
Distribution of SWI/SNF CRC components in MSC treatment groups and implications towards MSC chromatin regulation. **(A)** Venn diagram plot showing strictly shared and unique peaks between treatment groups. **(B)** Volcano plots showing significant peaks in red. Significant peaks defined as: average normalized counts>10 and FC>abs(2). 1 replicate per treatment group was analyzed. GSK343 treatment decreases the total number of accessible peaks and shows a lower number of upregulated DARs than downregulated DARs. **(C)** ARID1A and ACTL6A RNA-seq normalized counts plotted across treatment groups. **(D)** Visual summary of the epigenetic regulation of serial passaging and EZH2i role in blocking this effect.

ACTL6A is another core subunit of the SWI/SNF CRC that’s known to play part in the alternate SWI/SNF complex (esBAF) that can antagonize ARID1A-containing complexes. An overall increase in ACTL6A through serial passaging was observed (Fig. 5C), as RNA-seq data suggests. This increase in ACTL6A may cause a remodeling of the SWI/SNF complex leading to an eviction or impairment of the ARID1A subunit [32, 33]. This happens without a change in total ARID1A expression levels which is also evident from the RNA-seq gene expression level of ARID1A across treatments (Fig. 5C). This altered chromatin accessibility at key stemness genes could indicate a shift in the expression of a pro-stemness transcriptome, as shown later. As found in the ATAC seq motif analysis, TEAD2 was shown to be significantly upregulated in the GSK343 treated group indicating EZH2 inhibition, through preventing H3K27me3 deposition, YAP, sensitive to actin polymerization and mechanical stress, can now partner with TEAD transcription factors and bind to the accessible TEAD-binding sites. The YAP/TEAD complex has been shown to drive the transcription of genes that reinforce the stem-like state [34]. By lowering the ACTL6A, GSK343 could be indirectly promoting the reformation of functional, ARID1A-containing SWI/SNF complexes. This restored ARID1A-SWI/SNF complex can then actively remodel chromatin to maintain an open, transcriptionally permissive state, further reinforcing the stemness gene program initiated by YAP/TEAD and preventing the silencing of TEAD by PRC2. Lamin A was also stained for to understand if the mechanosensitive nuclear envelope protein shows any spatial localization in the MSC, but any Lamin A ring-like structure was not found in either early or later passage cells or upon any drug treatment (Fig. S3).

Overall, a model whereby serial passaging on a stiff substrate promotes loss of MSC stemness through a mechano-epigenetic pathway is proposed in Fig. 5D. Stiffness driven actin polymerization upregulates ACTL6A, which incorporates into the SWI/SNF complex, leading to the functional disruption of ARID1A-containing complexes. This impairs chromatin remodeling at stemness loci, allowing for their progressive silencing by EZH2-mediated H3K27me3. By inhibiting EZH2 with GSK343, H3K27me3 repressive mark is reduced, thereby breaking the cycle and promoting chromatin opening at key regulatory elements, such as TEAD2. This then facilitates the activity of mechanosensitive transcription factors like YAP/TAZ, which in tun downregulate ACTL6A. The subsequent re-establishment of functional ARID1A-SWI/SNF complexes creates a positive feedback loop that maintains an open chromatin landscape, thereby preserving the stemness of MSCs even at higher passages. This model provides a novel rationale for targeting epigenetic machinery to enhance the scalability and efficacy of MSC-based therapies.

### RNA sequencing reveals significantly differentially expressed genes between epigenetically modified MSC and early to late passaged MSC on a mechanically soft or stiff substrate

Next, our goal was to investigate the effect of stiffness induced chromatin architecture and its epigenetic mediation by drugs on the MSC gene expression. RNA sequencing was performed to characterize the change in the transcriptome of MSCs from serial passaging on the mechanically different substrates and epigenetic modifying drugs with each sample group. RNA sequencing results exhibited significant variability between samples as displayed in the principal component analysis (PCA) plot (Fig. 6). Technical replicates for each treatment group clustered together indicating low variance between replicates and a more similar transcriptome profile. Additionally, untreated P3 MSC groups (Control P3 and Soft P3) clustered together and late passage untreated P5 groups (Control P5 and Soft P5) clustered together. This data suggests that although the soft substrate is generally believed to maintain better MSC phenotype, as shown in our immunofluorescence study (Fig. 1, Fig. 2) and previous literature [35, 36, 18], the global gene expression level in late passage soft substrate primed MSC is not much different than the late passage MSC cultured on plastic. The GSK126 treatment showed the greatest amount of variability, indicating a large difference in their transcriptomics.

**Figure 6.**
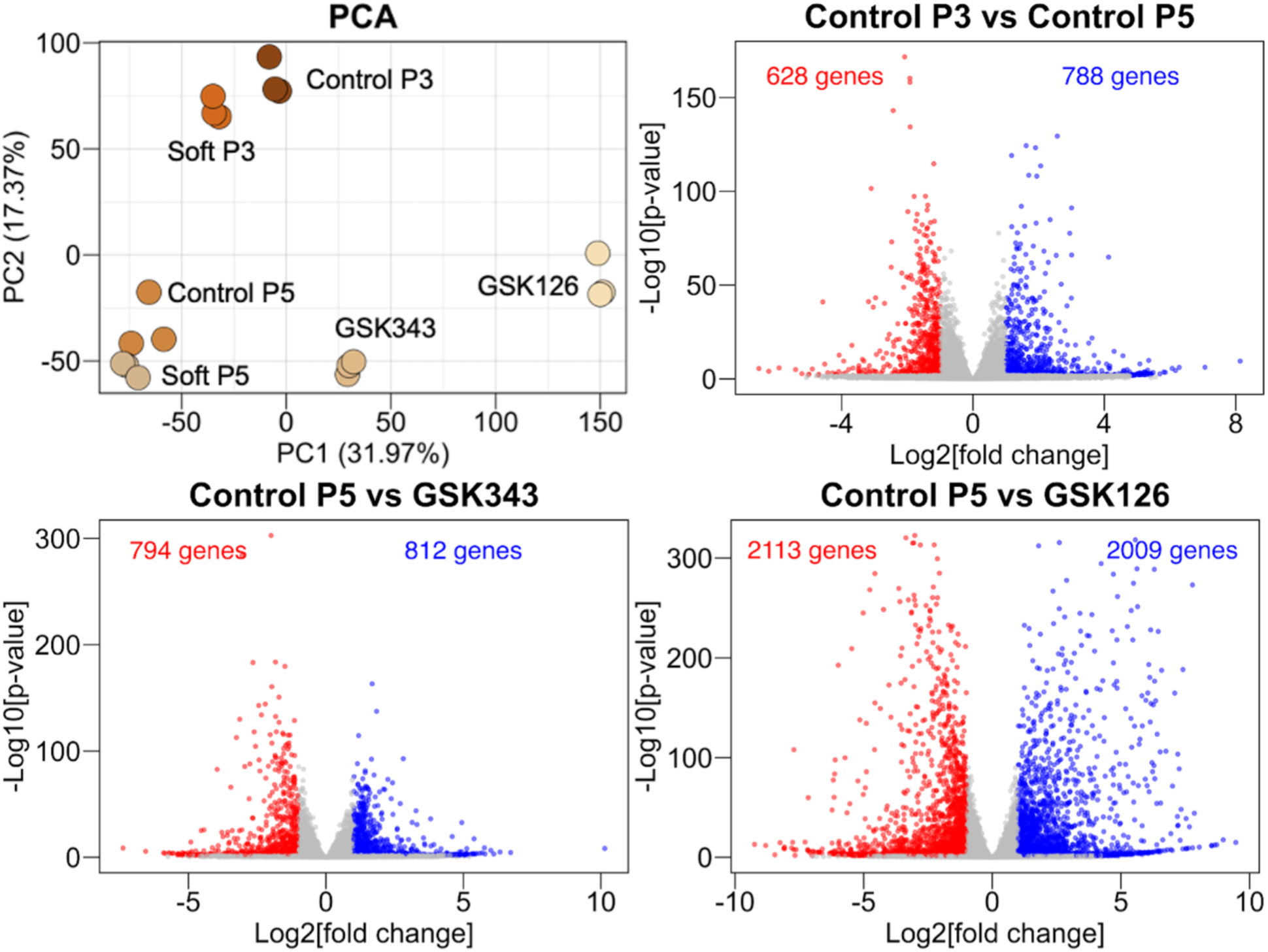
Stiffness dependent passaging and epigenetic modifications regulate the transcriptome of MSC. (A) PCA plot showing sample clustering based on gene expression profiles. About 50% of the total variance explained between PC1 (32%) and PC2 (17%). **(B)** Volcano plots displaying differentially expressed genes (DEGs). Significantly upregulated genes (FC > 1, FDR < 0.05) colored red and significantly downregulated genes (FC < -1, FDR < 0.05) are colored blue. A large variation in expression profiles was discovered between treatments.

A large number of differentially expressed genes (DEGs) were also illustrated in this data set. Analysis of the DEGs between groups showed many significantly up- or down-regulated genes. Out of the comparison groups analyzed, only Control P3-Soft P3 and Control P5-Soft P5 comparison groups showed less than 700 significant DEGs which correlates with the PCA plot variability (Fig. S4). Notably, GSK126 showed the largest number of significant DEGs compared to Control P5 further indicating its sizeable change in expression profile (Fig. 6). GSK343 also showed a large amount of DEGs in comparison to both Control P5 and Soft P5 groups. For Control P5-GSK343 comparison, which is the most relevant comparison from a translational perspective, 2194 genes were differentially expressed and of those, 1137 were up-regulated and 1057 genes were down-regulated. Overall, the initial analysis of the transcriptomic data showed a significant difference in gene expression between EZH2 inhibitors and cells cultured on varying culture substrates with different stiffnesses.

### Up and down- regulated DEGs indicate divergent transcriptome profiles in MSC treated with EZH2 inhibitors and early and late passage MSC

Upon further investigation of MSC associated genes (Fig. 7A), a heatmap depicting Z-score, illustrated functionally distinct phenotypes between the treatment groups. Normalized counts of specific genes of interest are shown in Fig. S5. Upregulation in genes such as CD34, FOXC1, and LEPR among others in the P3 untreated groups indicate a pro-inflammatory and undifferentiated state with high stemness. Conversely, the P5 untreated groups show upregulation in genes that indicate a hyperproliferative state. GSK126 pre-treatment showed upregulation in genes such as CDKN1A, CDKN2A, and IL6 among others that induce senescence-associated secretory phenotype (SASP) and anti-osteogenic effects. This hints that GSK126 pre-treatment induces senescence and potentially limits their therapeutic utility. GSK343 pre-treatment transcriptomic profile among MSC-specific genes shows a push toward an angiogenic phenotype with potential adipogenic priming (ANGPT1, SPP1, IGFBP7). Upregulation of genes such as ZEB2 and THY1 also suggest an increase in EMT and migration genes suggesting MSC plasticity and migration in an undifferentiated state.

**Figure 7.**
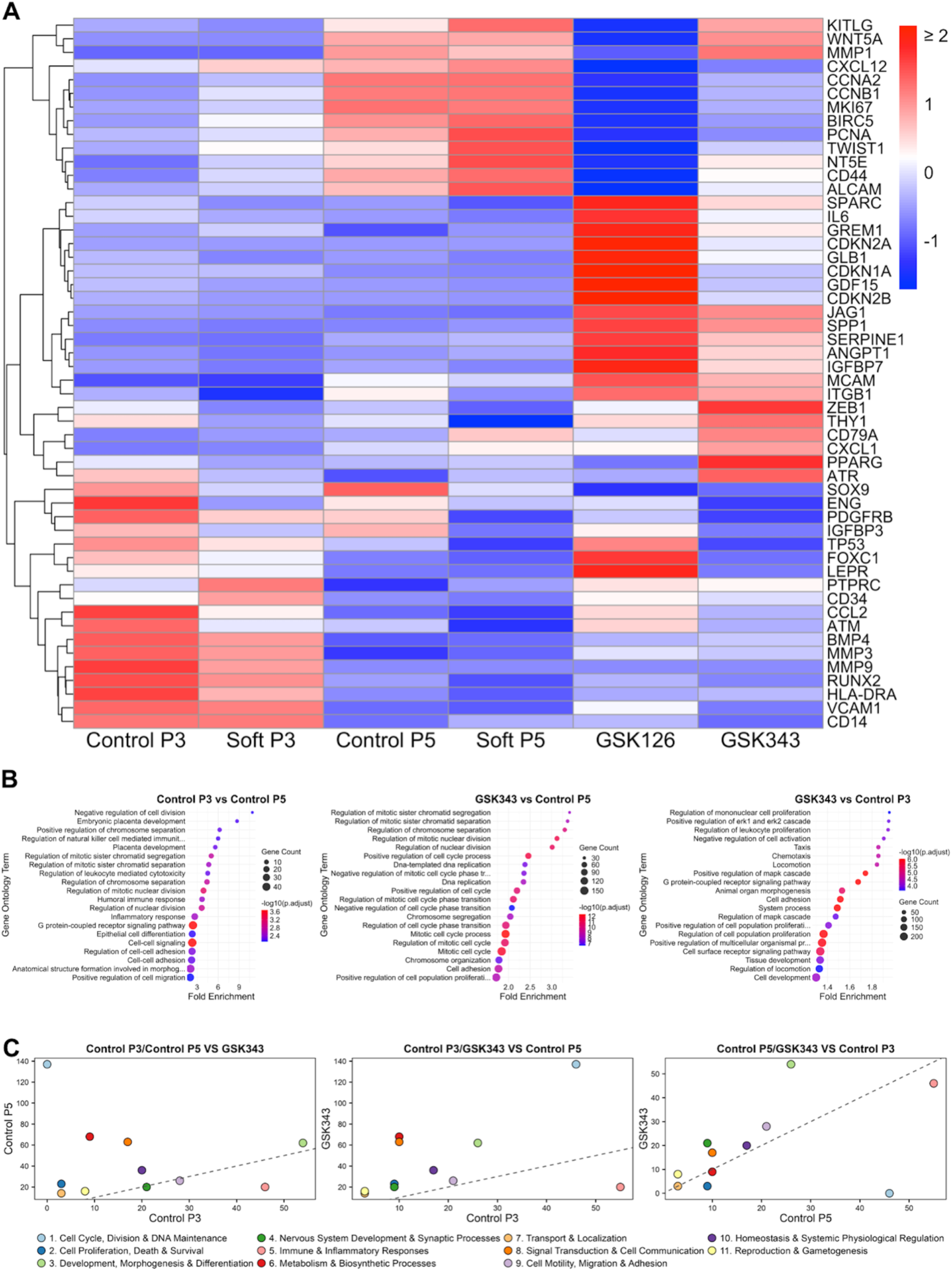
Assessment of substrate stiffness and epigenetic modification on MSC transcriptome. **(A)** Heatmap of normalized counts illustrating the Z-scores for MSC-specific genes in each treatment group. Red indicating increased expression and blue showing decreased expression compared to the mean. **(B) ORA pathway analysis of significant DEGs comparing Control P3, Control P5, and GSK343 treatment groups.** Top 20 most significantly affected pathways based on p-adjust value for each comparison. **(C)** Comparison of significantly affected pathways (adjust p-value<0.05) sorted into general biological categories between different treatment groups. The total number of pathways counted per category was plotted for each comparison.

To compare how each treatment condition influences biological pathways, over-representation analysis (ORA) on the significant DEGs (FDR<0.05) was performed from the comparisons Control P3 vs Control P5, GSK343 vs Control P5, and GSK343 vs Control P3. The pathways were sorted based on adjusted p-values and the top 20 most significant were plotted in Fig. 7B. In comparing early (P3) and late (P5) passage control groups, most of the top significantly affected pathways were related to cell division, DNA maintenance, and proliferation. There were also a few pathways associated with immunomodulatory and cell signaling processes. In comparing GSK343 to Control P5 a similar trend was observed where the top significantly affected pathways all had to do with cell cycle, DNA maintenance, and proliferation. Conversely, in comparing GSK343 to Control P3 only a few top pathways were related to proliferation. Instead, the most significant pathways were more diverse with pathways associated with immunomodulation, cell signaling, development, motility and adhesion.

To further investigate significantly affected pathways by the various treatments, all significant pathways (adjust p-value<0.05) were assigned to functional categories based on their biological roles. The number of pathways belonging to each category for each comparison was then quantified. To visualize this data, three scatterplots in which the x- and y-axes represent the pathway category counts from two contrasts at a time (Fig. 7C). Therefore, categories closer to the linear line illustrated on each plot indicate pathways that are similarly impacted by both conditions relative to the third group. Conversely, points that deviate strongly from the linear line identify categories in which one comparison shows greater pathway involvement than the other. This allows for a direct comparison of how pathway categories are altered between two treatment groups while highlighting shared and unique biological responses.

In comparing Control P3 and P5 to GSK343 a high divergence in the cell cycle, division, and DNA maintenance (1) category was observed indicating a lot of pathways were significantly affected in the Control P5 group compared to GSK343 where the Control P3 group had a more similar transcriptome profile to GSK343 in this category. Additionally, signal transduction and cell communication (8) and metabolism and biosynthetic processes (6) showed greater bias toward Control P5. On the other hand, the immune and inflammatory response (5) was shifted toward Control P3. When examining the shifts observed in Control P3 and GSK343 to Control P5 most categories were found to be more strongly enriched in the GSK343 group. This indicates an overall greater shift in the transcriptomic profile between GSK343 and Control P5 than between Control P3 and Control P5. The one group that was found to be more affected by the Control P3 was the immune and inflammatory response pathway. Lastly, in assessing how Control P5 and GSK343 groups differed from Control P3 the greatest shifts were found in the development, morphogenesis, and differentiation (3) category toward GSK343 and the cell cycle, division, and DNA maintenance (1) category toward Control P5.

### Proteomic analysis of the MSC conditioned media suggests soft substrate seeding and EZH2 inhibition promotes a more functional phenotype with implications towards fibrosis

The functional output of MSCs is defined by their secretome, which informs how cells communicate, their response to various stimuli, ECM remodeling, cell damage, and their metabolic activity. To examine these aspects, mass spectrometry of conditioned media from the treatment groups (Control P5, Soft P5, GSK343, and GSK126) was performed. Since only one replicate per group was tested, proteins were designated as significant if the |FC| was greater than 2 compared to the control media. The most significant proteins are listed in the heatmap in Fig. 8A and proteins that were uniquely found in some treatment groups are listed in the Venn diagram in Fig. 8B.

**Figure 8.**
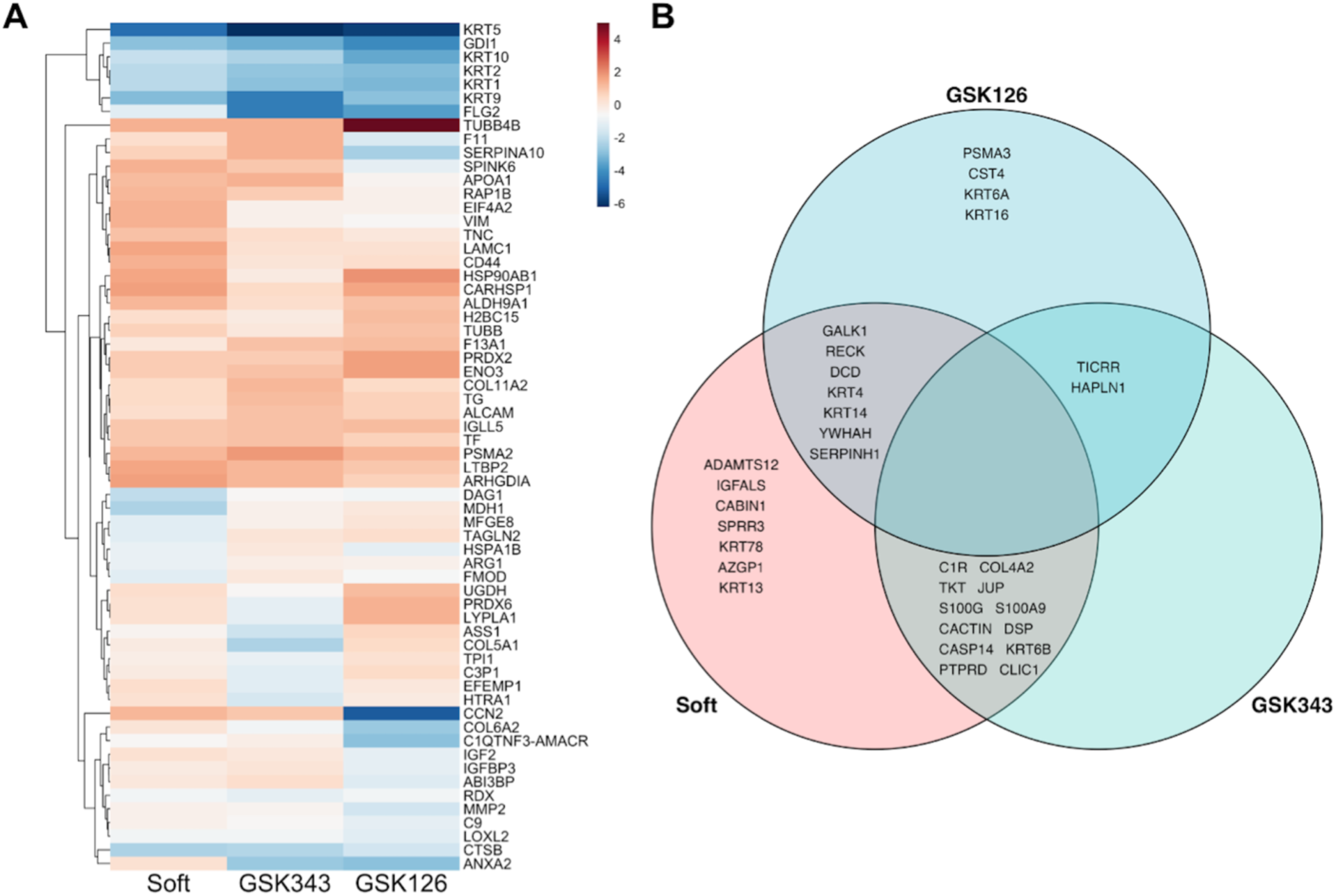
Mass spectrometry analysis of proteins in MSC media. **(A)** Heatmap of protein relative abundance values illustrating the log2(FC) for each MSC treatment group compared to the control. Red indicating up- regulation and blue indicating down-regulation of protein expression. **(B)** Venn diagram showing the proteins expressed by some treatment groups and not found in relative abundance in others.

For all treatment groups, a considerable downregulation in keratin proteins and FLG2 was observed. Keratins and FLG2 are hallmarks of epithelial cells indicating that MSC treatment potentially reduces the epithelial-like transition in MSC and promotes a more primitive multipotent state. Additionally, all groups showed an increase in TUBB4B relative expression compared to the control which could point to enhanced MSC motility and paracrine signaling.

GSK126 also shows a reduction in genes related to ECM (COL6A2, MMP2, LOXL2, etc) aligning with EZH2’s role in fibrosis. However, GSK126 treated cells had an upregulation of PRDXs and HSP90AB1 which could suggest a compensatory stress mechanism where GSK343 showed downregulation of these proteins. Similarly, GSK343 also reduces collagen production such as COL4A2 and COL5A1 suggesting an altered matrix stiffness response and stronger ECM modulation (Fig. 8B). Protein expression diverges further between the two EZH2i treatments when GSK343 uniquely upregulated SERPINA10 and F11. This suggests that GSK343 treatment might aid in modulating immune responses. The softer substrate stiffness seems to be mimicking the effects of EZH2 inhibition in keratin suppression; however, ADAMTS12 and CLIC1 expression are upregulated drastically indicating a divergence in phenotype around ECM remodeling.

Further investigating secreted proteins unique to specific treatments, PSMA3 was upregulated in GSK126 compared to the Control CM by a log2(FC) of 4.2 and RECK was downregulated by log2(FC) of -2. These changes could indicate a loss in MSC ability to remodel their ECM or reduced matrix degradation and may drive pro-inflammatory secretory profiles. Conversely, ADAMTS12 expression was highly upregulated in the soft treatment media indicating an increase in the MSC ability to degrade their ECM and promote tissue flexibility. GSK343 and soft treatment share 12 proteins unique to them. Notably, JUP was downregulated by over log2FC of -4 by both groups indicating a loss of epithelial traits in the MSC. S100A9 was also downregulated by both groups and especially GSK343 treated cells by log2FC of -4. S100A9 has been shown to promote senescence in MSC therefore downregulation of S100A9 could prevent senescence during passaging.

### Conditioned media from cultured MSCs and EZH2 inhibition increase chondrocyte proliferation in vitro

The MSC secretome has a wide range of soluble mediators that have been shown to play a significant role in repairing and modulating tissues [37]. More specifically, previous studies show that the MSC secretome has beneficial effects on cultured chondrocytes by promoting their proliferative, migratory, and secretory capacities [38]. To further elucidate how EZH2 inhibition effects MSC therapeutics through their secretome MSC conditioned media (CM) was collected, and chondrocytes were cultured in MSC CM from MSCs that either received GSK343 treatment (GSK343 CM) or non-treated MSC (Control CM). The chondrocytes were given a 1:1 or 3:1 ratio of basal media to MSyC CM. Control wells consisted of the same ratios but with MSC media instead of MSC CM. As expected, chondrocytes that received MSC conditioned media in any ratio were found to increase chondrocyte proliferation compared to the control wells (Fig. 9). Additionally, EZH2 inhibition by GSK343 increased the chondrocyte proliferation with a significant difference shown in the 75:25 groups. Interestingly, the 75:25 ratio of basal media to CM was found to increase proliferation compared to their 50:50 counterparts. Generally, it appears that EZH2 inhibition through GSK343 enhances the MSC secretory profile that can benefit chondrocyte proliferation.

**Figure 9.**
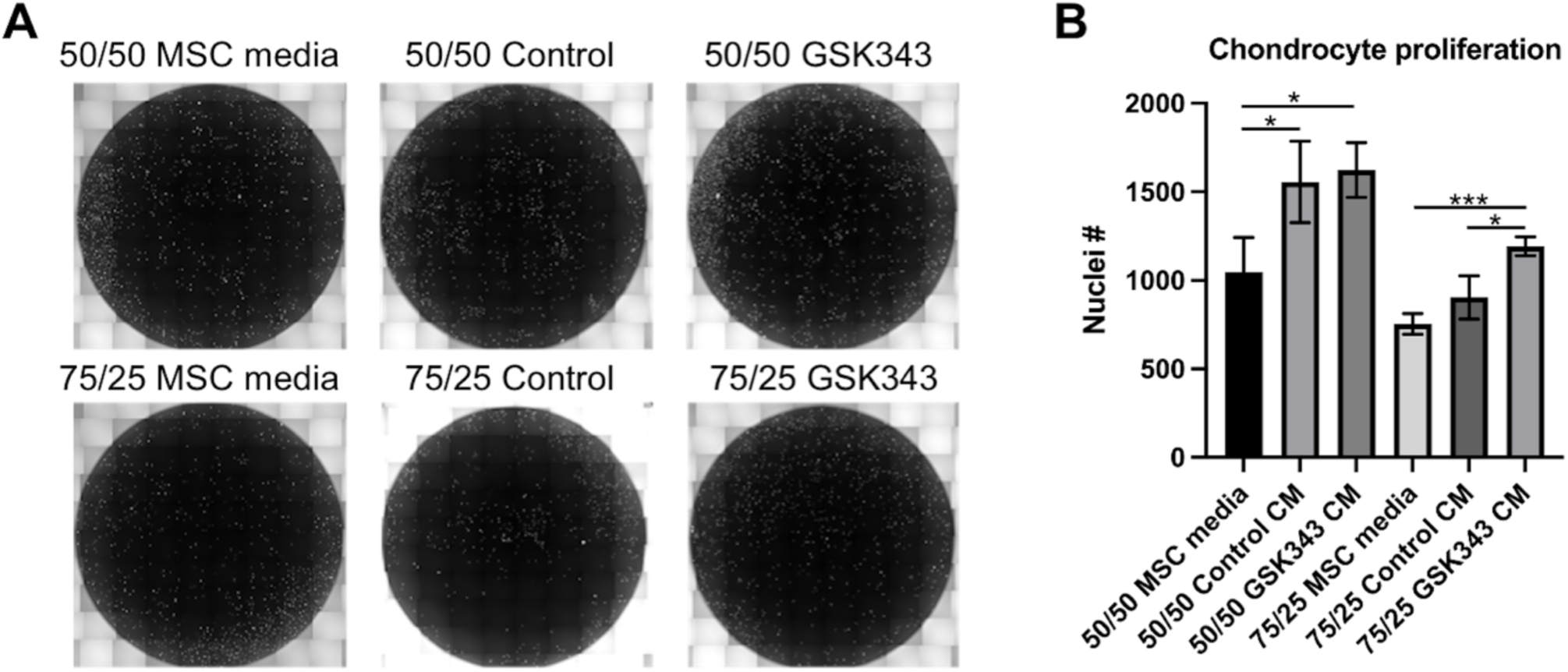
Chondrocyte proliferation assay with MSC-conditioned media (CM). **(A)** Representative 10x tiled images of whole well stained with DAPI for each treatment group. **(B)** Cell count from DAPI stained nuclei on day 4 post seeding cells at same density. **Conditioned media from GSK343 treated cells promotes greater chondrocyte proliferation.**

## CONCLUSION

This study demonstrates that selective EZH2 inhibition -particularly with GSK343 - can effectively preserve mesenchymal stem cell (MSC) stemness during serial passaging on stiff, non-physiological substrates by recapitulating key mechano-epigenetic features normally maintained on soft, bone marrow-like environments. Across phenotypic, epigenetic, chromatin-accessibility, transcriptomic, and proteomic analyses, GSK343 consistently promoted a stem/progenitor-like state, sustained expression of critical MSC markers, and partially restored chromatin architecture disrupted by passage-induced mechanical stress. Mechanistically, our findings support a model in which GSK343 interrupts stiffness-driven PRC2–SWI/SNF imbalance, reduces H3K27me3- mediated silencing, and re-enables TEAD/YAP-associated regulatory networks that underpin MSC stemness. Functionally, GSK343 reshaped the MSC secretome toward a more therapeutic profile and enhanced chondrocyte proliferation, underscoring its potential relevance for regenerative applications such as cartilage repair and inflammatory/fibrotic conditions.

Future work should validate these findings in vivo and across MSCs from multiple donors to assess reproducibility and therapeutic durability. Longitudinal studies combining GSK343 with engineered mechanical microenvironments may further optimize expansion strategies for clinical-scale MSC manufacturing. Additionally, dissecting the direct genomic targets of GSK343-mediated chromatin opening - particularly TEAD/YAP-regulated loci - and mapping how SWI/SNF subunits dynamically redistribute during passaging will deepen mechanistic insight. Finally, evaluating the therapeutic impact of GSK343-conditioned MSC secretome in disease-relevant models of osteoarthritis, fibrosis, or immune dysregulation could establish a foundation for next-generation epigenetically primed MSC therapies.

## CONFLICT OF INTEREST

The authors declare no conflict of interest

## ACKNOWLEDGEMENT

The authors acknowledge grant funding from NIH/ NCATS Colorado CTSA UL1 TR002535 and NSF CAREER 2236710. The authors acknowledge Natalie Calahan, Kanita Hrustanovic, Scott Burlingham, Kenna Thomas, Emily Kaplan, Jack Forman, Sawyer Halingstad and Ryan Mahoney for their contributions in experiments and data analysis. The authors acknowledge the funding (NSF Grant 2117943 and RRID SCR 021758) and staff from the Bioanalysis and Omics core facility at CSU.

## SUPPLEMENTARY MATERIALS

**Figure S1.**
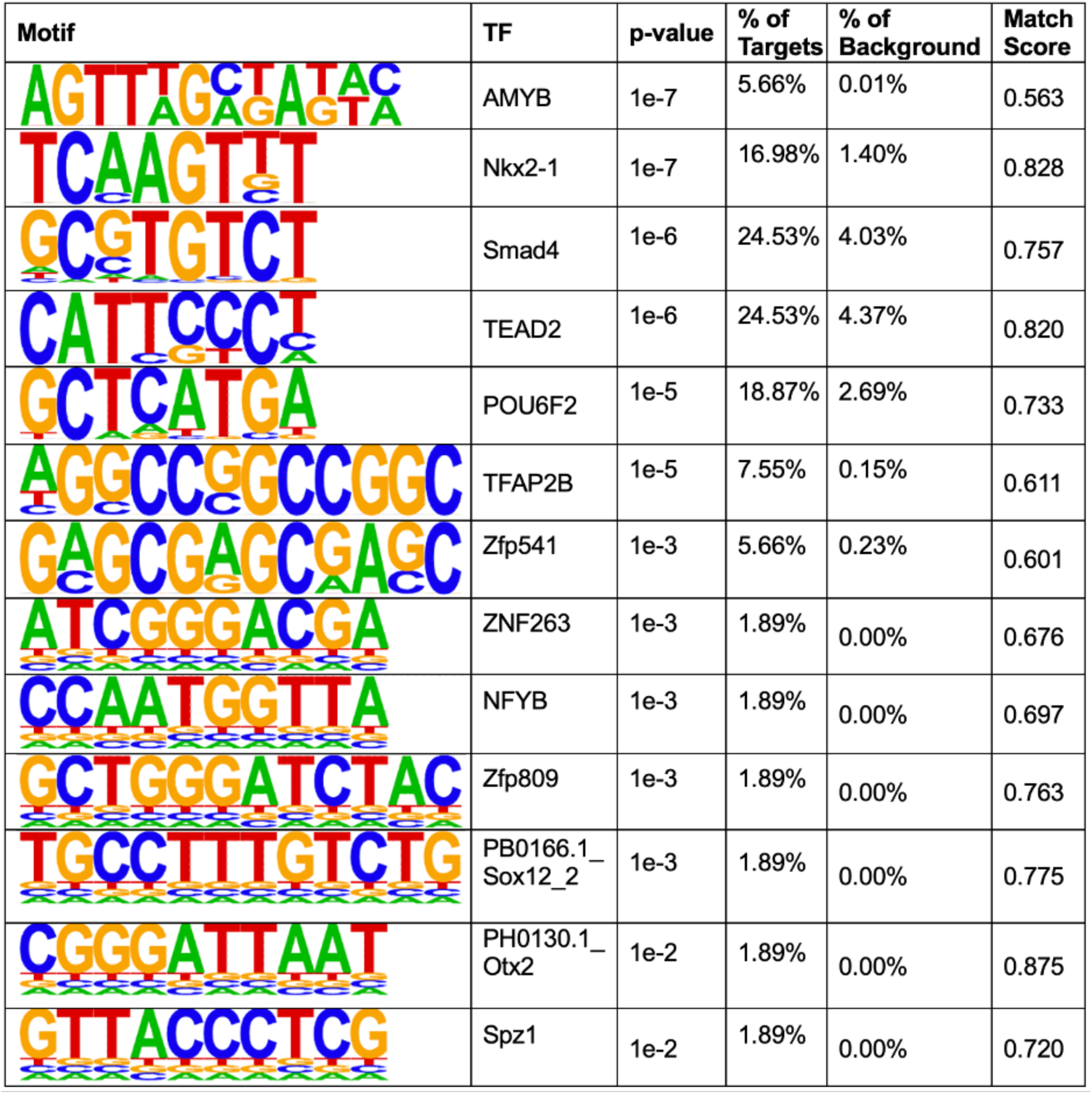
Homer motif analysis of the 53 significantly upregulated peaks in GSK343 vs Control.

**Figure S2.**
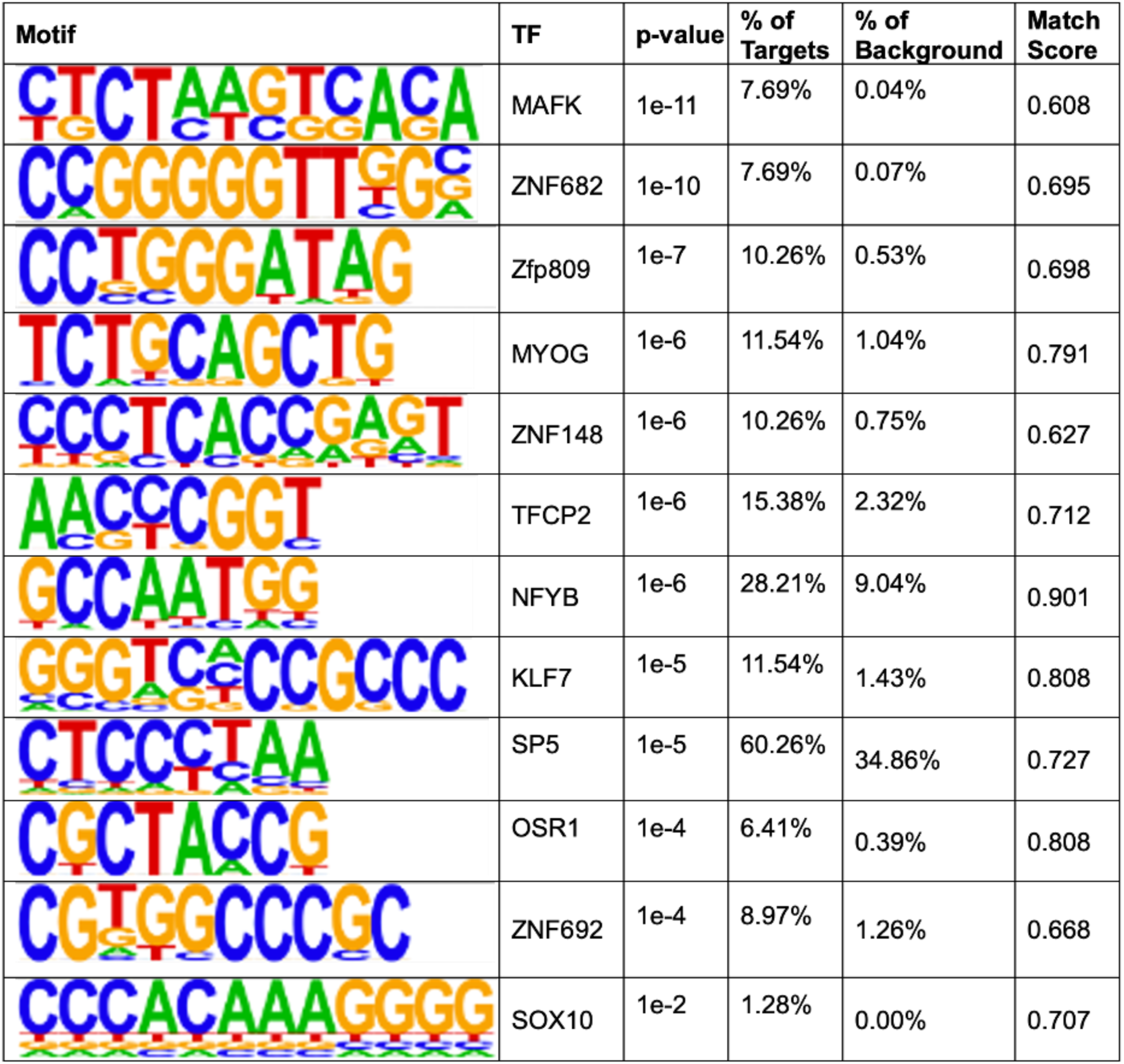
Homer motif analysis of the 78 significantly downregulated peaks in GSK343 vs Control.

**Figure S3.**
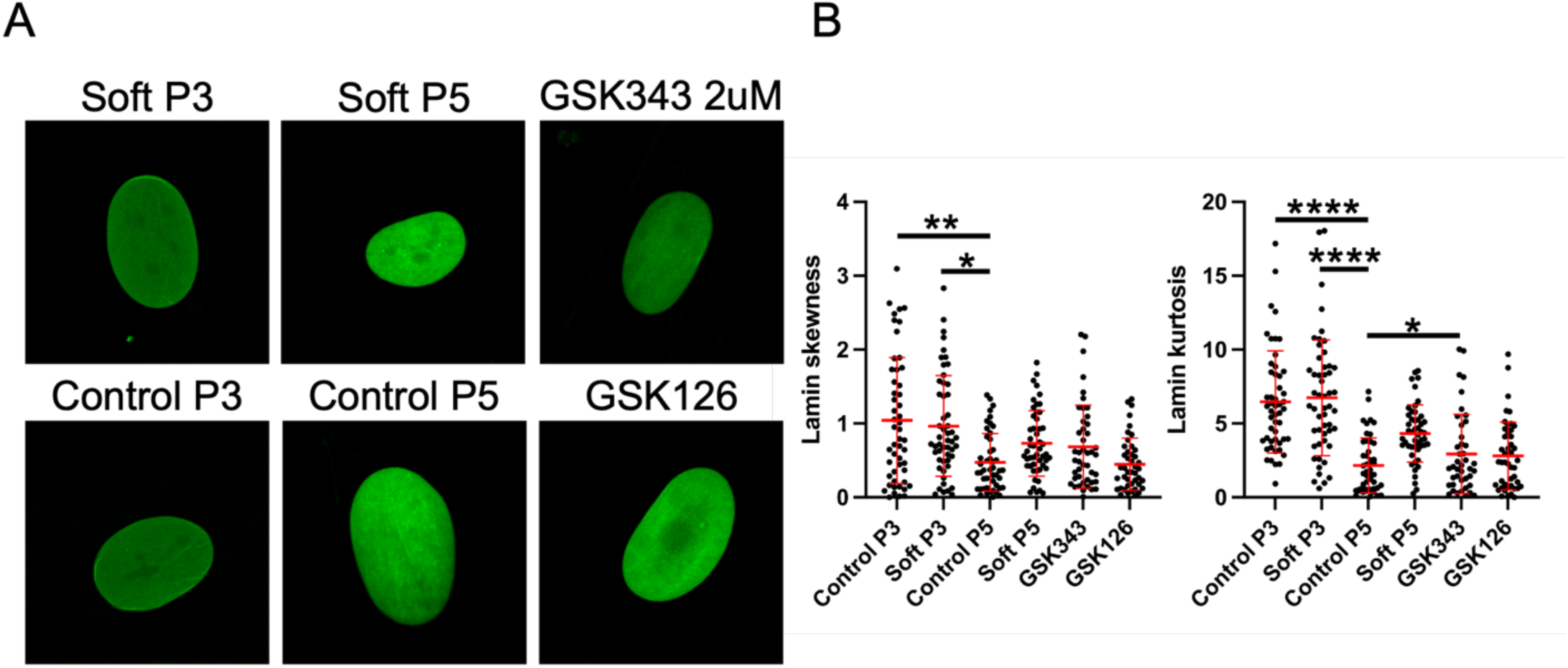
Spatial distribution of Lamin A protein is shown in MSC. No specific spatial pattern was visible in either early or later passage cells not it was affected by any of the drugs.

**Figure S4.**
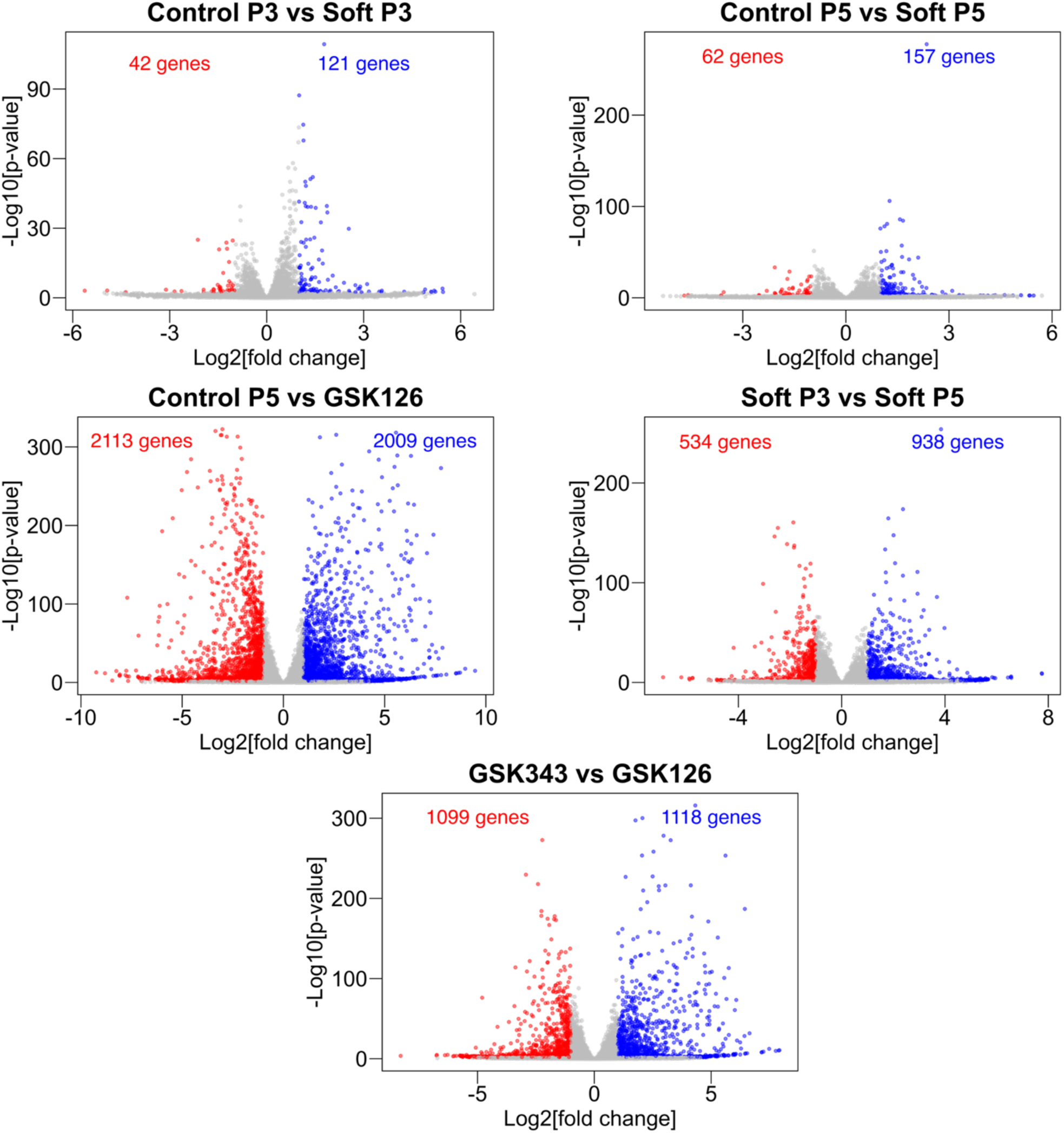
Volcano plots for several pairwise comparisons.

**Figure S5.**
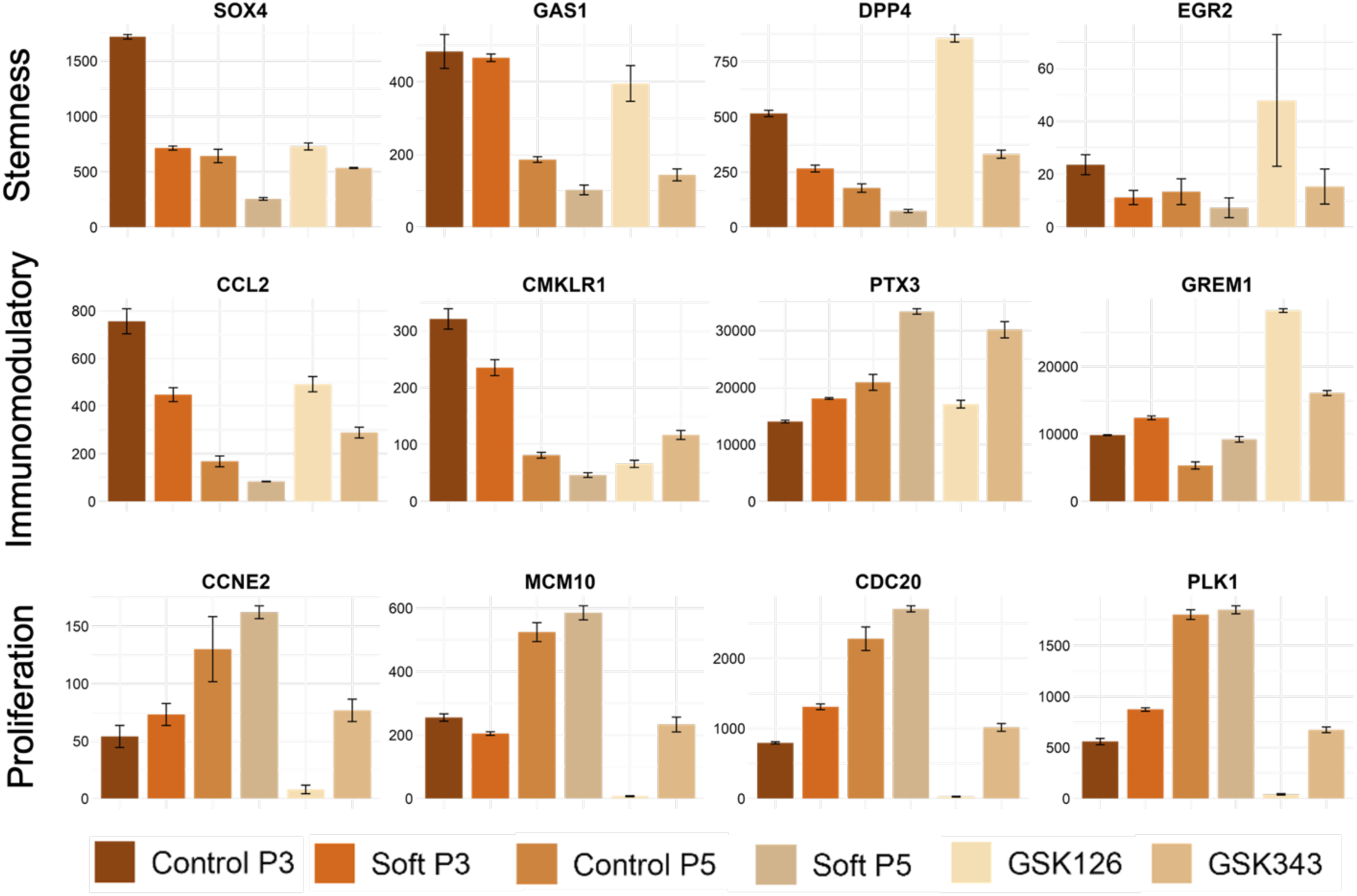
Plots for normalized counts of specific genes derived from transcriptomics data.

## Notes

### Competing Interest Statement

The authors have declared no competing interest.

